# Point-of-care peptide hormone production enabled by cell-free protein synthesis

**DOI:** 10.1101/2022.12.03.518932

**Authors:** Madison A. DeWinter, Ariel Helms Thames, Laura Guerrero, Weston Kightlinger, Ashty S. Karim, Michael C. Jewett

## Abstract

In resource-limited settings, it can be difficult to safely deliver sensitive biologic medicines to patients due to cold chain and infrastructure constraints. Point-of-care drug manufacturing could circumvent these challenges since medicines could be produced locally and used on-demand. Towards this vision, we combine cell-free protein synthesis (CFPS) and a 2-in-1 affinity purification and enzymatic cleavage scheme to develop a platform for point-of-care drug manufacturing. As a model, we use this platform to synthesize a panel of peptide hormones, an important class of medications that can be used to treat a wide variety of diseases including diabetes, osteoporosis, and growth disorders. With this approach, temperature-stable lyophilized CFPS reaction components can be rehydrated with DNA encoding a SUMOylated peptide hormone of interest when needed. Strep-Tactin^®^ affinity purification and on-bead SUMO protease cleavage yields peptide hormones in their native form that are recognized by ELISA antibodies and that can bind their respective receptors. With further development to ensure proper biologic activity and patient safety, we envision this platform could be used to manufacture valuable peptide hormone drugs at the point-of-care in resource-limited settings.

## Introduction

Peptide hormones are a class of biologic drugs with a range of therapeutic benefits, particularly in the treatment of chronic diseases, which disproportionately affect people living in resource-limited settings(*1*). Diseases like diabetes(*2*), obesity (*3,4*), and osteoporosis(*5*) are commonly treated with peptide hormone biologics. However, current centralized manufacturing processes for biologics present challenges, highlighted by the COVID-19 pandemic, in both the transportation and storage of sensitive biologic materials.

Packaging to minimize mechanical stresses and maintenance of cold chain temperatures, which require both reliable energy sources and temperature monitors, are important considerations for the appropriate handling of sensitive biologic therapeutics(*6*). The high cost and need for reliable infrastructure of these measures makes it difficult to deliver these drugs safely and effectively to resource-limited settings. By nature, it is hard to accurately predict *a priori* the quantity of a drug that will be needed. Therefore, there tends to either be a shortage or an excess of each desired drug with current centralized manufacturing processes. When there is excess drug product that cannot be used before its expiration, it must be disposed of properly or risk contaminating water supplies(*7*), which adds an additional cost burden. Taken together, the current approach for biomanufacturing peptide and protein therapeutics can limit response time, distribution of medicines, and drive healthcare inequities. This backdrop sets the stage to reimagine drug manufacturing away from current reliance on centralized facilities (*6,8,9*).

Cell-free protein synthesis (CFPS) has recently emerged as one potential way to rethink decentralized drug manufacturing. CFPS is a technology that harvests the transcription and translation machinery from cells to produce peptides and proteins *in vitro*(*10–12*). Over the last two decades, yields of CFPS systems have increased to more than a gram per liter(*13–15*), system costs have been reduced(*16–18*), and glycosylation, a key feature of many therapeutics, has been enabled(*19–26*). Towards decentralized manufacturing, CFPS is advantageous because it can be lyophilized for transportation and storage at ambient conditions(*27,28*) thus eliminating the need for maintenance of cold-chain storage. After transportation and storage, CFPS reactions can be rehydrated on-demand at the point-of-care to synthesize the drug of interest(*23,29*).

Previous work in CFPS systems has demonstrated synthesis of a protective *Francisella tularensis* glycoconjugate vaccine(*23*), an enterotoxigenic *Escherichia coli* (ETEC) vaccine(*27*), antimicrobial peptides, a diphtheria vaccine, nanobodies, and DARPins(*29*), granulocyte colony stimulating factor (*30,31*), and onconase(*32*), among others. While these efforts have covered numerous therapeutics, peptide biologics have been underrepresented due to challenges in CFPS production. Protease degradation makes it difficult to produce small peptides in crude-extract based CFPS and recombinantly in *E. coli*(*33–35*).

Here, we developed a platform for the cell-free production of peptide biologics in crude extracts. First, we developed and optimized a strategy to increase expression and stabilize a panel of peptide hormones in CFPS. A key feature was the use of fusion proteins to help improve expression and purification of peptides (*35,36*). Specifically, we selected a small ubiquitin-related modifier (SUMO) protein fusion to help stabilize the expressed peptide hormones and to provide a handle for an on-bead, simultaneous enzymatic cleavage and purification system. Second, we developed a simultaneous 2-in-1 enzymatic purification scheme that isolates unmodified peptide hormones. Finally, we demonstrated peptide activity through receptor binding. Overall, this work represents a first step to express peptide hormones in crude extract-based CFPS systems for decentralized manufacturing.

## Results and Discussion

We sought to develop a workflow that would allow for point-of-care peptide hormone production by CFPS (**Figure 1**). The key concept would be to ship pre-made, lyophilized CFPS reactions at ambient temperatures to local manufacturing sites where the medication is *needed*(*23,29*). The lyophilized CFPS reactions, which are stable at ambient temperatures(*27*), can then be rehydrated as needed with the plasmid DNA encoding the needed peptide hormone. After expression, the hormones would be purified for delivery to patients.

**Figure 1:**
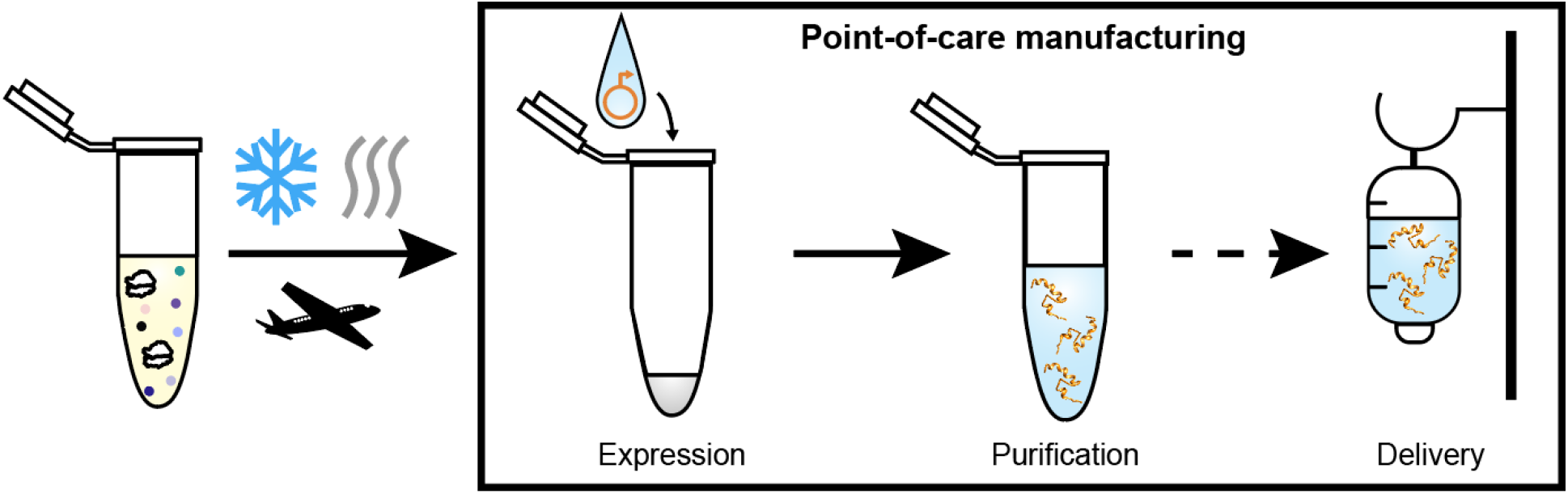
Lyophilized cell-free expression enables point-of-care peptide hormone manufacturing, purification, and patient delivery. A proposed scheme for point-of-care manufacturing where centrally manufactured CFPS reactions are lyophilized for transportation and storage at room temperature. When needed, the lyophilized CFPS pellets can be rehydrated with plasmid encoding the desired peptide hormone. After the CFPS reaction is complete, the peptide hormones can be purified to eliminate background lysate material and delivered to patients.

To develop this platform, we selected twenty peptide hormones ranging from 1 kDa to 22 kDa as model peptides based on their potential for expression in cell-free systems (e.g., minimal or no disulfide bonding, no non-canonical amino acids, no post-translation modifications) and their therapeutic use (**Supplementary Table 1**). For example, glucagon-like peptide 1 (GLP-1) receptor agonists, like semaglutide and liraglutide, have been used for the treatment of diabetes(*37*) and more recently, have been highlighted for their use in obesity(*38,39*). Growth hormone is used for both congenital growth hormone deficiencies in children and deficiencies secondary to systemic diseases like chronic kidney disease and cancer(*40–42*). Parathyroid hormone can be used in the treatment of both osteoporosis(*43*) and hypoparathyroidism(*44*).

### Peptide hormones can be expressed as soluble biologics in point-of-care enabled CFPS

We first attempted to express each of the twenty selected hormones in standard BL21 Star™ (DE3) extract using CFPS. After CFPS, we only observed full-length expression of 4 out of the 20 tested hormones (**Figure S1a**). Consistent with previous observations(*33,34*), successfully expressed peptide hormones (insulin A and B chains fused with leucine zipper heterodimers(*45*), growth hormone, and leptin) tended to be the largest (>10 kDa) of the peptide hormones in the panel. The larger bands in the insulin A and B chain heterodimer lanes may be due to unintended dimerization or trimerization.

To increase expression and stabilize the peptides during CFPS, we employed a fusion protein *strategy*(46,47). We decided to use small ubiquitin-related modifier (SUMO) protein as the fusion protein for its ability to improve molar expression of recombinantly expressed proteins(*46,48*) and to be cleaved using a protease (ubiquitin-like protease-1, Ulp-1) that leaves a scarless product(*46,48–50*). Indeed, the addition of the SUMO tag to the peptide hormones improves expression of peptide hormones in CFPS using standard crude BL21 Star (DE3) lysate (**Figure S1b**). With the SUMO fusion, 8 of the 20 peptide hormones in the panel were expressed at full-length, showing a clear improvement over the untagged hormones. There are bands below the peptides of interest that migrate similarly to where SUMO protein (12 kDa) would be, indicating that there may be truncations and/or non-specific protease cleavage.

We then investigated whether changing the protein content of our CFPS reactions, by using crude extracts derived from different *E. coli* strains, improved SUMOylated peptide hormone expression (**Figure S2**). To establish a baseline, we employed the minimal PUREfrex 2.1 synthesis system and successfully expressed all 20 SUMOylated peptide hormones at full-length (**Figure S2b**). Although purified cell-free synthesis systems are stable after lyophilization at room temperature for at least 1 year(*51*), PUREfrex 2.1 is expensive and has significantly lower yield after lyophilization when compared to lyophilized crude extracts(*52*), making it poor for scale up. We then tested an *E. coli* K strain engineered for *in vitro* glycosylation, Clm24 (W3310 Δwaa)(*53*), that has separately been engineered to have a modified lipopolysaccharide structure to reduce its endotoxin effects(*23*). The modified lipopolysaccharide could simplify downstream purification and processing of therapeutics generated using our workflow. We found that 14 of the 20 peptide hormones expressed at full-length using Clm24 (**Figure S2c**), offering an improvement over the BL21 lysate. Finally, we tested a release factor 1 (RF-1)-deficient K strain of *E. coli* containing a proteolytic-resistant T7 RNA polymerase on the genome, 759.T7.Opt, that creates some of the highest-yielding extracts reported(*14*). The 759.T7.Opt strain performed the best, synthesizing 17 out of the 20 peptide hormones (**Figure S2d and S2e**), and was selected for the following experiments.

With an improved expression system consisting of 759.T7.Opt extracts and SUMOylated hormone constructs, we performed an initial purification screen to produce soluble peptide hormones (**Figure S3**). The entire panel of peptide hormones was subjected to affinity purification and cleavage of the SUMO protein to leave untagged peptide hormones. The GLP-1 mutant, GLP-1, growth hormone (GH), and parathyroid hormone (PTH) were successfully recovered as detected by a Coomassie-stained SDS-PAGE gel.

We moved forward with these four peptide hormones and then quantified their expression using radioactive ^14^C-leucine incorporation during CFPS. We found that each of our SUMOylated hormones expresses at about 20-60 μM, higher than the untagged counterparts (**Figure 2a**). An autoradiogram confirms that the SUMOylated hormones express as full-length constructs (purple arrows, **Figure 2b**), with smaller bands likely resulting from truncations and/or non-specific protease cleavage.

**Figure 2:**
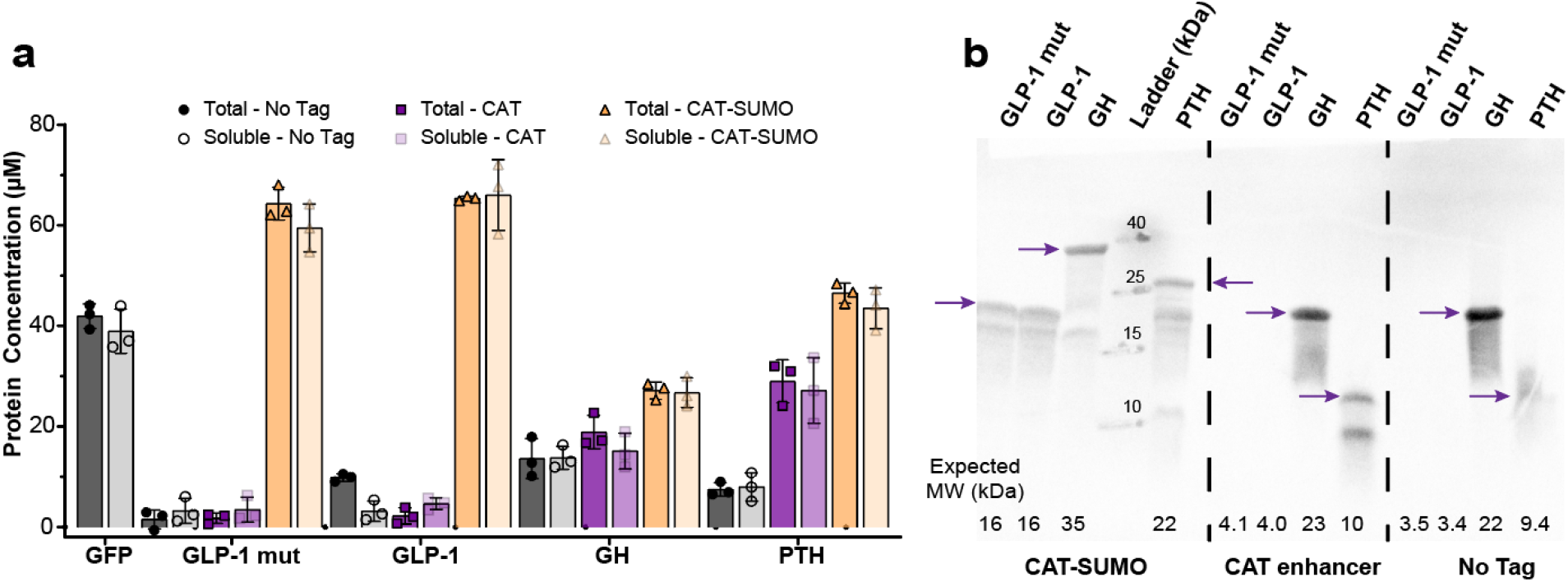
Quantitative determination of the improvement of SUMO fusion peptide hormone expression over untagged hormones. (a) Incorporation of radioactive ^14^C-leucine was used to quantify protein expression of untagged (black), N-terminal CAT-enhancer sequence tagged (purple) (**Supplementary Table 2**), and CAT-enhancer SUMO fusion (orange) hormones produced in CFPS. Darker shades represent total hormone expression and lighter shades represent soluble expression. GFP was included as a well-expressing, soluble positive control for comparison. N=3 replicates were performed for each sample, and error bars represent 1 s.d. (b) An autoradiogram of an SDS-PAGE gel of a mixture of the same samples from (a) shows that full-length constructs are expressed (purple arrows). Hormones with a CAT-enhancer SUMO fusion are shown on the left, with a CAT-enhancer only in the middle, and without any tags on the right.

To improve access to sensitive biologic drugs like these peptide hormones in resource-limited settings, CFPS production would require lyophilization of the reactions, eliminating the need to maintain cold chain during transportation and storage (*23,29*). Therefore, we assessed peptide production after lyophilization. With the exception of GH, lyophilization does not significantly impact soluble yields of CFPS produced hormones. Yields of SUMOylated GLP-1 mut, GLP-1, and PTH synthesized in lyophilized CFPS were statistically equivalent to those of hormones produced in CPFS reactions that had not been lyophilized (“standard”) (**Figure 3a**). An autoradiogram was performed to confirm the full-length constructs were expressed (**Figure 3b**). As with “standard” expression (**Figure S2**), lyophilized reactions contain both full-length peptide (purple arrows) and protein that runs at approximately the molecular weight of SUMO.

**Figure 3:**
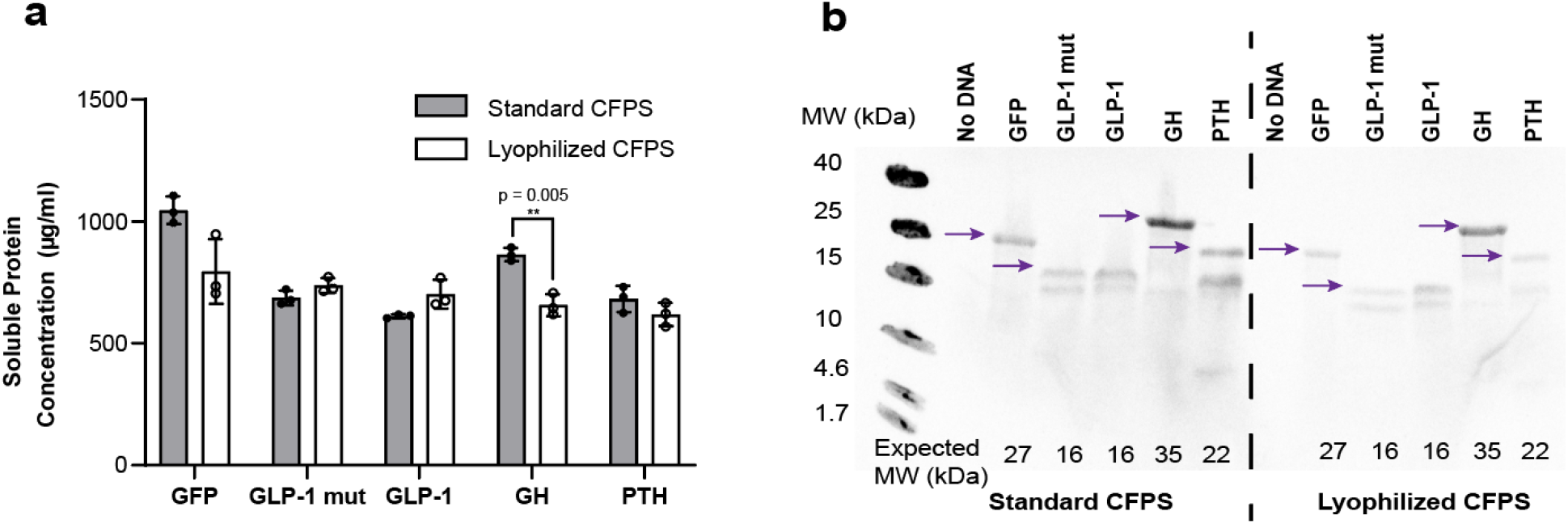
Fusion with a SUMO protein allows successful expression of peptide hormones in both standard and lyophilized CFPS reactions. (a) Quantification of StrepII-SUMO-hormone production by radioactive ^14^C-leucine incorporation is represented for both standard (dark grey) and lyophilized (white) CFPS reactions. All samples (n=3) are shown with error bars representing 1 s.d. Samples were compared using two-tailed t-test assuming samples with unequal variance. (b) An autoradiogram of an SDS-PAGE gel for a mixture of the replicates from panel (a) is shown. Purple arrows point to expected masses.

### A simultaneous 2-in-1 enzymatic purification scheme isolates unmodified peptide hormones

We next set out to design a scheme to both purify the peptide hormones from the background crude lysate and to cleave the SUMO protein off the synthesized hormones to leave untagged, and presumably active, hormones. Our 2-in-1 purification scheme involved binding the N-terminal StrepII-SUMO peptide hormone fusion to Strep-Tactin^®^ functionalized magnetic beads and washing away background CFPS contaminants (**Figure 4a**). Ulp-1 protease then recognizes the tertiary structure of SUMO and cleaves precisely at the SUMO C-terminus to release untagged hormone(*50*). Because the StrepII tag remains at the N-terminus of SUMO, SUMO remains bound to the magnetic beads. Nickle chelate beads are added to remove the His-tagged Ulp-1 protease from solution and the purified, untagged peptide hormone is collected.

**Figure 4:**
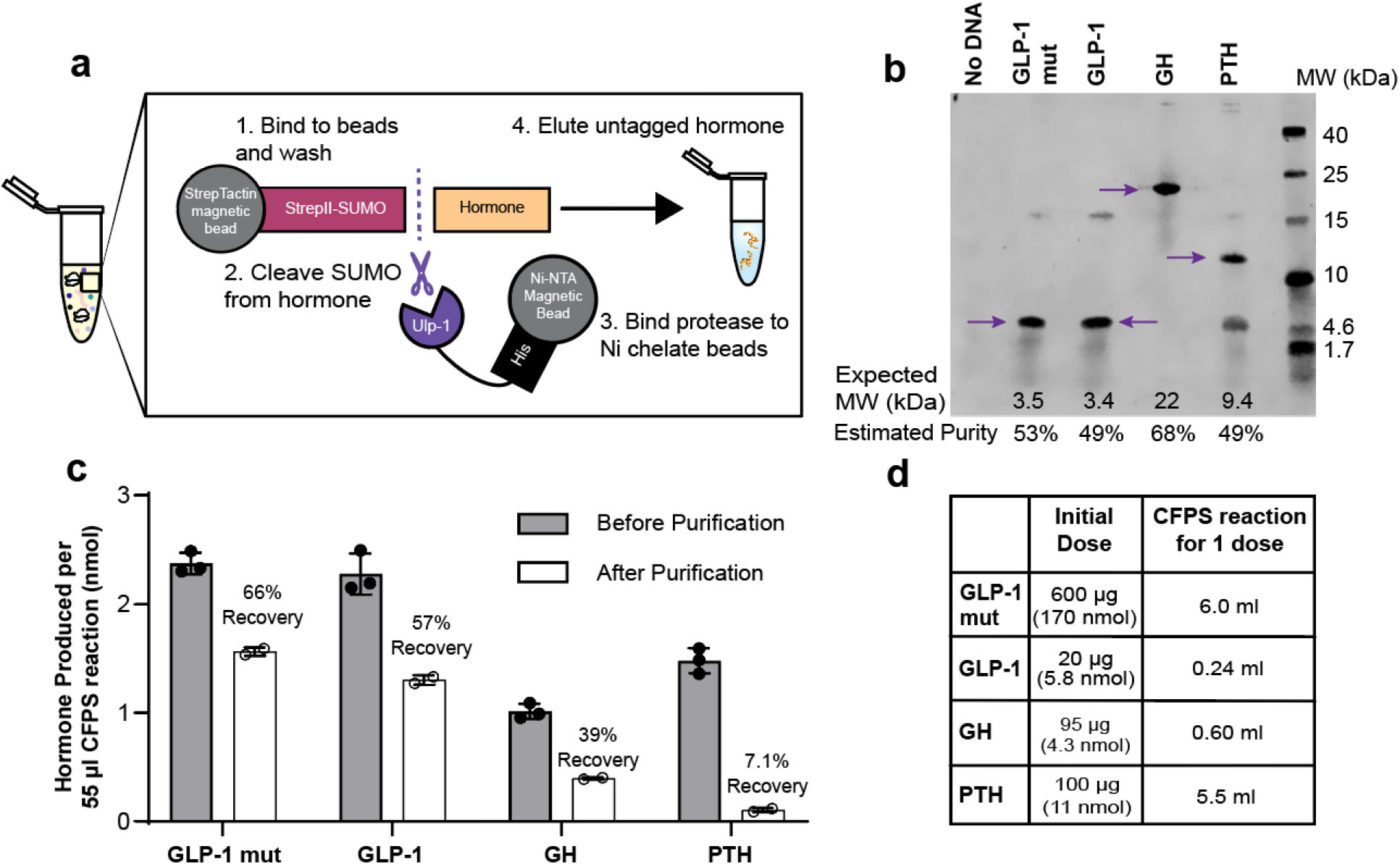
A simultaneous 2-in-1 enzymatic purification scheme isolates unmodified peptide hormones. (a) The scheme for 2-in-1 purification is shown. (b) A Coomassie stained SDS-PAGE gel shows intact, tagless peptide hormones after purification. The band in each lane corresponding to the full-length, tagless hormone is highlighted with a purple arrow. Densitometry was used to obtain the estimated purity. (c) A BCA assay was used to quantify the total protein recovery after purification (n=2) from a 55 μl CFPS reaction and the hormone recovery was calculated based on the purity by densitometry in panel (b) (white). The recovery was calculated by comparing to the production measured by radioactive leucine incorporation before purification (dark grey) (n=3). Error bars represent 1 s.d. (d) A table showing the typical initial dose for each of these medications and the CFPS reaction volume necessary to produce the dose assuming the same yields from panel c are maintained during scale up.

Previous work has been done where enzymatic cleavage reactions were used to elute protein off purification resin with a TEV protease variant(*54*), a picornavirus 3C protease(*55*), a PreScission™ protease(*56*), and even with scSUMO(*57*). However, to our knowledge, this work represents the only effort to use SUMO to both stabilize the expression of therapeutics and to provide a handle for purification of a scarless version of the peptide or protein. We used this purification scheme to purify the peptide hormones to approximately 50% purity by densitometry on a Coomassie-stained SDS-PAGE gel (**Figure 4b**). The amount of hormone recovered varied greatly, ranging from 7.1% to 66%, between each hormone (**Figure 4c**). This yield is low compared to typical affinity purification schemes; however, it theoretically could be improved by further optimizing SUMO cleavage conditions or changing the method of affinity purification from a magnetic bead scheme to a set up with resin in a column.

We used the purification yields to calculate the size of CFPS reactions that would be necessary to produce a therapeutic dose (**Figure 4d**). We determined that GLP-1 mutant would require 6.0 ml of CFPS to produce enough GLP-1 mutant for 1 dose (assuming a typical liraglutide dose of 600 μg(*58*)). Typically, GLP-1 receptor agonists have modifications that improve half-life; however, GLP-1 has been used in research to treat patients with type 1 diabetes using a continuous infusion of 1.2 pmol/kg/min of GLP-1 throughout a meal(*59*). Assuming an 80 kg adult that eats for 60 minutes, this would correspond to a total of 20 μg (5.8 nmol). Producing this amount of GLP-1 would require a 240 μl CFPS reaction. Growth hormone is typically dosed by weight and 0.33 mg/kg/week is a normal initial dose(*41*). For an average 20 kg 5-year-old boy, the daily dose would be 95 μg or 600 μl of CFPS reaction. To treat osteoporosis, a typical initial dose of parathyroid hormone is 100 μg(*60*) or 5.5 ml of CFPS reaction. Given CFPS reactions have been shown to scale up to 100 L(*61*), the CFPS reaction volumes required for one dose of any of those therapeutics (~0.6-6ml) is well within the range of feasibility.

### Characterization by LC-MS, ELISA, AlphaLISA, and BLI shows cell-free produced growth hormone is intact and capable of binding its receptor

The size and function of the purified peptides are important to verify for therapeutic uses. We next sought to confirm the cleaved hormones are the appropriate size by coupled liquid chromatography-mass spectrometry (LC-MS), ensure recognition of the hormones by two orthogonal antibodies in a sandwich enzyme-linked immunosorbent assay (ELISA), and test hormone binding to its respective receptor with both an amplified luminescent proximity homogeneous assay (AlphaLISA) and biolayer interferometry (BLI).

We first showed this for growth hormone (**Figure 5**). Protein LC-MS after treatment of the sample with dithiothreitol (DTT) revealed an observed mass of 22,129.34 Da from maximum entropy deconvolution which matches the theoretical GH mass (22,129.05 Da) with an error of 13 ppm (**Figure 5a**). GH was also recognized by two orthogonal antibodies in a commercial sandwich ELISA and showed a recovery of 72 ± 7 μg/ml (**Figure 5a**). This is approximately 45% of the amount detected by BCA and densitometry. We also assessed GH engagement with the extracellular domain of its receptor with AlphaLISA (**Figure 5b**). AlphaLISA is an assay that detects protein-protein interactions based on the proximity of proprietary donor and acceptor beads. GH was expressed with an sFLAG tag(*62*) on the C-terminus to immobilize to anti-FLAG donor beads. The commercially produced extracellular domain of the growth hormone receptor (GHR, Acro GHR-H52222) was His-tagged and immobilized on nickel chelate acceptor beads. A cross-titration of both growth hormone and its receptor revealed a pattern consistent with clear binding, suggesting that the GH produced in CFPS and purified using our workflow can recognize and engage with its receptor (**Figure 5b**). The binding was confirmed with BLI (**Figure 5c**), an orthogonal method to optically detect macromolecular interactions. Although a K_D_ value for the hormone-receptor binding was not able to be determined because there was no significant dissociation observed, clear binding was observed during association (**Figure 5c**). The observed association on BLI is consistent with the receptor engagement observed by AlphaLISA in **Figure 5b**.

**Figure 5:**
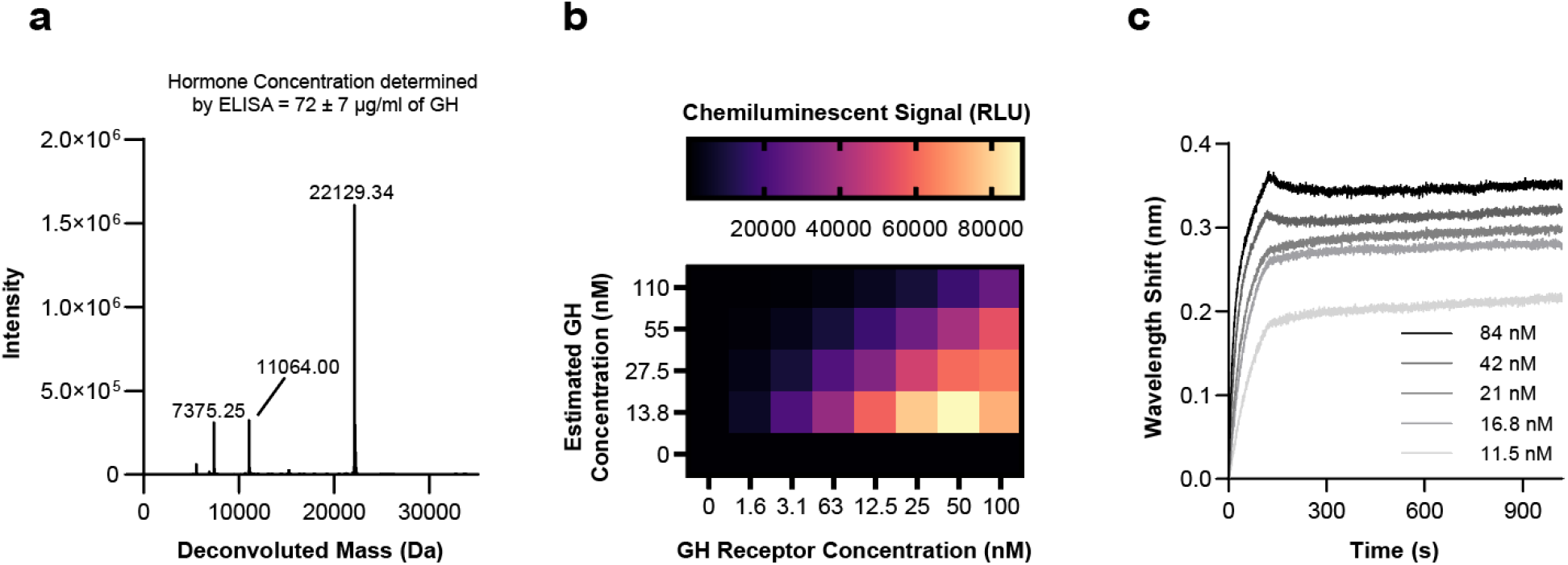
Characterization of GH by ELISA, LC-MS, AlphaLISA, and BLI shows cell-free produced growth hormone is intact and capable of binding its receptor. (a) GH identification and quantification was determined by sandwich ELISA with 4 dilutions and intact protein LC-MS. The observed mass (22129.34 Da) is shown on a deconvoluted spectrum, and it matches the expected average reduced mass of GH (theoretical: 22129.05 Da) with an error of 13 ppm. (b) GH and GH receptor concentrations were cross titrated, and binding was assessed with AlphaLISA. (c) GH binding to the extracellular domain of its receptor was assessed by BLI. Kinetic curve fitting was unable to be performed due to lack of observed dissociation. Data are representative of at least three independent experiments.

We also set out to characterize GLP-1, its mutant (**Figure S4**), and PTH (**Figure S5**). GLP-1 and the GLP-1 mutant were detected as full-length peptides in peptide LC-MS (**Figure S4a-b**). The peptide hormones are cleaved precisely at the C-terminus of SUMO to give tagless hormones as expected. Using sFLAG tagged GLP-1, anti-FLAG donor beads, Fc-tagged extracellular domain of the GLP-1 receptor, and Protein A acceptor beads, AlphaLISA detected GLP-1 binding to its receptor 3 standard deviations above background (**Figure S4c**). Unfortunately, this binding was unable to be confirmed with BLI, likely because the hormone is too small (3.5 kDa) to generate enough of a wavelength shift to be detected by the instrument (**Figure S4d**). Full-length parathyroid hormone was detected in the deconvoluted LC-MS spectra and via sandwich ELISA (**Figure S5a**). The concentration was determined to be 24 ± 11 μg/ml (**Figure S5a**). This value is approximately 80% of that detected by BCA and densitometry. AlphaLISA suggested that PTH interacts with the extracellular domain of its receptor, but signal was low (**Figure S5b**). Unfortunately, due to high background binding and a relatively small PTH (9.5 kDa), BLI was also unable to detect PTH binding to the extracellular domain of its receptor (**Figure S5c**).

We then wanted to extend this expression and purification method to a panel of antimicrobial peptides. After the full expression and purification scheme, we were able to recover five peptides detected by Coomassie stained SDS-PAGE – Casocidin-II, Melittin, Bactericidin B-2, Catestatin, Thrombocidin-1 D42K, and Thrombocidin-1 (purple arrows, **Figure S6a**). In an overnight incubation of *E. coli* strain MG1655 in a 384 well plate in two replicates, melittin was shown to be functional by inhibiting growth of the bacteria (**Figure S6b**), demonstrating that this workflow can be used in applications outside of peptide hormones.

Overall, we created a cell-free expression and first-pass purification procedure for peptide hormone production to be performed at the point-of-care. Further development efforts are necessary to ensure that the hormones would be safe for injection into patients, particularly with regards to the purity of the final product. Endotoxin removal and polishing purification steps would also need to be integrated into the system. However, this work represents a step towards the ability to reproducibly synthesize biologics, including peptide hormones, in a decentralized approach in resource-limited settings. This work also expands the synthesis capabilities of CFPS for therapeutically relevant molecules, particularly for biologics with molecular weights smaller than 10 kDa.

## Materials and Methods

### Genes

All constructs were expressed in the pJL1 plasmid (Addgene 69496). A CAT enhancer sequence (MEKKI) was present at the N-terminus, followed by a linker (RGS) to the Strep-II tag (WSHPQFEKTG). The SUMO protein follows without a linker and then the peptide hormone is at the C-terminus of the SUMO protein. For antimicrobial peptides, the gene is in the same location as the peptide hormone. The amino acid sequences for these constructs are given in **Supplementary Table 2**. For sFLAG constructs designed for AlphaLISA experiments, a GS7 linker (GGGSGGG) separates the sFLAG amino acid sequence evolved for higher affinity to anti-FLAG antibodies (DYKDEDLL)(*62*) from the C-terminus of the hormone. The heterodimer(*45*) is located after a GSSGS linker at the C-terminus of the affinity tag.

### Preparation of cell extracts for CFPS

Crude extracts for CFPS were made from several *Escherichia coli* strains. The “CLM24” strain is the W3110 strain with a knockout of the ligase that transfers the O polysaccharide to the lipid-A core (WaaL), engineered for N-linked protein glycosylation(*53*). The “BL21” strain is the BL21 Star™ (DE3) (Fisher Scientific) strain, and the “759” strain has a release factor 1 (RF1) knockout from the C321.ΔA strain with integration of T7 RNA polymerase with mutations K183G and K190L to prevent OmpT cleavage onto the genome (759.T7.Opt)(*14*). Cells were grown in 2xYTP media (5 g/L sodium chloride, 16 g/L tryptone, 10 g/L yeast extract, 7 g/L potassium phosphate monobasic, and 3 g/L potassium phosphate dibasic, pH 7.2) in 1 L shake flasks or in a Sartorius Stedim BIOSTAT Cplus 10 L bioreactor. BL21 and CLM24 were grown at 37 °C and 759 was grown at 34 °C. At an OD_600_ of ~ 0.6, T7 RNA polymerase expression was induced with 1 mM IPTG in the BL21 and 759 strains. At an OD_600_ of ~2.8, cells were harvested and pelleted by centrifugation at 8,000 xg for 5 min at 4 °C. Cells were washed with chilled S30 buffer (10 mM Tris-acetate pH 8.2, 14 mM magnesium acetate, 60 mM potassium acetate) 3 times and pelleted by centrifugation at 10,000 xg for 2 min at 4 °C. 1 mL/g wet cell mass of S30 buffer was used to resuspend BL21 and CLM24 cells and 0.8 ml/g wet cell mass of S30 buffer was used to resuspend 759 cells. 1 ml aliquots of BL21 and 1.4 ml aliquots of 759 cells were lysed by a Q125 Sonicator (Qsonica) at 50% amplitude. BL21 cells were pulsed 10 s on, 10 s off until 640 J were reached, and 759 cells were pulsed for 45 s on and 59 s off until 950 J were reached. CLM24 cells were lysed with single pass through an Avestin EmulsiFlex-B15 at 20,000 to 25,000 psig. The lysates were centrifuged for 10 min at 4 °C. BL21 was spun at 10,000 xg and 759 and CLM24 were spun at 12,000 xg. The supernatant was collected and BL21 was aliquoted and flash frozen. 759 and CLM24 both underwent runoff reactions for 1 hr at 37 °C with 250 rpm shaking and were spun at 12,000 xg and 10,000 xg, respectively, for 10 min at 4 °C. Supernatants were collected and samples were aliquoted, flash frozen, and stored at −80°C until use.

### Cell-free protein synthesis (CFPS)

55 μl CFPS reactions were carried out in 2 ml microcentrifuge tubes (Axygen) using a modified PANOx-SP system(*18*). 10 mM magnesium glutamate, 10 mM ammonium glutamate, 130 mM potassium glutamate, 0.85 mM each of GTP, UTP, and CTP, 1.2 mM ATP, 34 μg/ml folinic acid, 0.171 mg/ml *E. coli* tRNA, 0.33 mM NAD, 0.27 mM CoA, 4 mM oxalic acid, 1 mM putrescine, 1.5 mM spermidine, 57 mM HEPES, 2 mM 20 amino acids, 0.03 M phosphoenolpyruvate, and 13.3 mg/ml plasmid purified using Zymo Midiprep Kits was added to each reaction. If lyophilized, 30 mg/ml of sucrose was added as a lyoprotectant. 45.8 μl of pre-mix with all components except for plasmid were dried for at least 4 hours after flash freezing on a VirTis BenchTop Pro lyophilizer (SP Scientific) at 100 mtorr and −80°C. Lyophilized pre-mix was rehydrated with 45.8 μl of nuclease free water (Fisher Scientific AM9937) and then added to tubes containing 9.2 μl of 80 mg/ml of plasmid DNA. Reactions were incubated overnight at 30 °C.

### Radioactive Quantification of CFPS Yields

Soluble CFPS yields were quantified by incorporation of ^14^C-Leucine (PerkinElmer) as previously described(*63*). After lyophilization, reactions were rehydrated with ^14^C-Leucine in water to give a final 10 μM concentration in triplicate 15 μl reactions. These reactions were transferred to tubes containing the appropriate plasmid and incubated overnight at 30 °C. 6 μl of soluble fractions were taken by removing the supernatant of reactions spun at 16.1k xg for 10 min at 4 °C and amino acids were cleaved off charged tRNA by incubation at 37 °C for 20 min with 0.25 N KOH. 5 μl of solutions were spotted on a 96 well filtermat (PerkinElmer 1450-421), dried, then precipitated by a 5% solution of trichloroacetic acid at 4 °C. The filtermat was dried and radioactivity was measured by a liquid scintillation counter (PerkinElmer MicroBeta) compared to a filtermat with the same solutions that had not been precipitated by trichloroacetic acid.

### Autoradiograms

Samples that were quantified by radioactive ^14^C-Leucine incorporation were run on an SDS-PAGE gel to confirm full-length expression. Samples were prepared for SDS-PAGE by adding 5 μl of sample to 4 μl of tricine sample buffer (Bio-Rad 1610739) and 1 μl of 1 M DTT and heat denaturing at 90 °C for 10 min. The samples were run on an 16.5% MINIPROTEAN^®^ Tris-Tricine gel (Bio-Rad 4563066) at 100 V for 90 min at constant voltage. The gels were dried (Hoefer GD2000) and exposed for at least 4 days before imaging on a Typhoon FLA 700 (GE Healthcare).

### Hormone Purification

After overnight CFPS expression, 35 μl of IBA MagStrep “type3” XT Strep-Tactin beads were equilibrated in a wash buffer of tris-buffered saline solution with 0.05% Tween 20 (ThermoFisher 28360) by washing twice with 35 ul of the wash buffer. Wash steps involve separating the magnetic beads from solution by using a magnetic plate and removing the supernatant before adding fresh wash buffer and resuspending the magnetic beads. 55 μl of wash buffer was then added to each CFPS sample and the solution was added to the magnetic beads and spun at ~18 rpm on a rotator at 4 °C for at least 45 minutes. The beads were then washed 3 times with 120 μl of wash buffer and resuspended in 30 μl of wash buffer. 5 μl of 1 U/μl of Ulp-protease (SigmaAldrich SAE0067) was added to each sample and incubated at room temperature with spinning at ~18 rpm for at least 2 hours. 2 μl of Dynabeads™ His-Tag Isolation and Pulldown beads (ThermoFisher 10104D) for each CFPS reaction were equilibrated twice in equal volumes of wash buffer before addition to the purified reactions to remove the Ulp-protease and incubated with spinning at 4 °C for at least 30 minutes. The purified solution was removed from the magnetic beads and placed in fresh tubes for further analysis.

### SDS-PAGE with Coomassie

Electrophoresis was used to separate peptides in a 16.5% Mini-PROTEAN^®^ Tris-Tricine Gel (Bio-Rad) with tris-tricine (Bio-Rad) or 4x Protein Sample Loading Buffer (LiCOR) and DTT. Tris-tricine running buffer (Bio-Rad 1610744) was used to run the gel at 100 V for 90 minutes. The gel was stained for at least an hour with Bulldog Bio 1-Step Coomassie stain (AS001000) and destained overnight in water.

### Densitometry

A built-in densitometry software (Image Studio Lite) was used to compare the brightness of the band of interest to the entire lane. The ratio of these values was used to determine the purity of the sample. The purity was calculating by taking the ratio of the intensity of the band of interest to the intensity of the entire lane. There are relatively clean bands (purple arrows) with minor contaminants in all the lanes containing peptide hormones. Because of the nature of the Ulp-1 protease cleavage site directly after the last two glycine residues at the C-terminus, these bands represent tagless hormones with a precisely defined amino acid sequence lacking any scar sequence or remaining amino acids resulting from the original fusion with SUMO.

### BCA

Total protein concentration in the purified samples was determined with the Pierce Rapid Gold BCA Protein Assay kit (ThermoFisher A53225) by comparing absorbance signal at 480 nm to a BSA standard curve ranging from 0-2000 μg/ml using manufacturer’s instructions. Hormone samples were run in duplicate with a 10 μl sample volume and 200 μl of working reagent in clear flat-bottomed 96 well plates (Corning 3370).

### Calculations to determine recovery after purification

After using a bicinchoninic acid (BCA) assay to quantitate total protein in the purified samples, purity from the densitometry analysis in **Figure 4b** was used to calculate the final peptide hormone concentration. The percent purity was multiplied by the total protein concentration to obtain the recovered, purified protein concentration.

### Protein LC-MS

Intact hormone molecular weight of large (>6 kDa) hormones was determined with LC-MS by injection of 5 μl of the purified peptide hormone from 3 separate reactions. After surfactant removal with the acidic protocol of the ProteoSpin detergent removal and sample concentration kit (Norgen Biotek 22800), samples were injected into a Bruker Elute UPLC equipped with an ACQUITY UPLC Peptide BEH C4 Column, 300 Å, 1.7 μm, 2.1 mm× 50 mm (186004495 Waters Corp.) with a 10 mm guard column of identical packing (186004495 Waters Corp.) coupled to an Impact-II UHR TOF Mass Spectrometer (Bruker Daltonics, Inc.). Liquid chromatography was performed using 0.1% formic acid in water as Solvent A and 0.1% formic acid in acetonitrile as Solvent B at a flow rate of 0.5 mL/min and a 50 °C column temperature. An initial condition of 20% B was held for 1 min before elution of the proteins of interest during a 4 min gradient from 20 to 50% B. The column was washed and equilibrated by 0.5 min at 71.4% B, 0.1 min gradient to 100% B, 2 min wash at 100% B, 0.1 min gradient to 20% B, and then a 2.2 min hold at 20% B, giving a total 10 min run time. An MS scan range of 100–3000 m/z with a spectral rate of 2 Hz was used. External calibration was performed. Data was analyzed using the maximum entropy deconvolution algorithm built-in to the Bruker DataAnalysis software.

### Peptide LC-MS

Peptide hormone molecular weight of small (<6 kDa) hormones was determined with LC-MS by injection of 5 μl of the purified peptide hormone from 3 separate reactions. After surfactant removal with the acidic protocol of the ProteoSpin detergent removal and sample concentration kit (Norgen Biotek 22800), samples were injected into a Bruker Elute UPLC equipped with an ACQUITY UPLC Peptide BEH C18 Column, 300 Å, 1.7 μm, 2.1 mm× 100 mm (186003686 Waters Corp.) with a 10 mm guard column of identical packing (186004629 Waters Corp.) coupled to an Impact-II UHR TOF Mass Spectrometer. Liquid chromatography was performed using 0.1% formic acid in water as Solvent A and 0.1% formic acid in acetonitrile as Solvent B at a flow rate of 0.5 mL/min and a 40 °C column temperature. An initial condition of 5% B was held for 1 min before elution of the proteins of interest during a 6.1 min gradient from 5 to 100% B. The column was washed for 2 min at 100% B and equilibrated at 5% B for 2.85 min, giving a total 12 min run time. An MS scan range of 100–3000 m/z with a spectral rate of 8 Hz was used. External calibration was performed.

### AlphaLISA

AlphaLISA assays use the proximity of donor and acceptor beads to detect protein-protein interactions. One protein is immobilized on donor beads that are excited by a laser to release a singlet oxygen species that can diffuse up to approximately 100 nm in solution. If the beads are in close proximity due to the interaction of the two species, the singlet oxygen will diffuse to the acceptor bead and trigger another chemical reaction to release a chemiluminescent signal that can be detected by a plate reader(*64*). AlphaLISA assays were run based on a previous protocol (*65*) in a 50 mM HEPES pH 7.4, 150 mM NaCl, 1 mg/ml BSA, and 0.015% v/v Triton X-100 buffer (“Alpha buffer”). An Echo 525 acoustic liquid handler was used to dispense materials from a 384-well Polypropylene 2.0 Plus source microplate (Labcyte, PPL-0200) using the 384PP_Plus_GPSA fluid type into a ProxiPlate-384 Plus (Perkin Elmer 6008280) destination plate. Anti-FLAG donor beads (Perkin Elmer) were used to immobilize the peptide hormone and either Protein A (Fc tagged GLP-1 receptor, Sino Biological, 13944-H02H) or Ni-NTA (6x His tagged GH, Acro GHR-H5222 and PTH, Abcam ab182670, receptors) acceptor beads (PerkinElmer) were used to immobilize the extracellular domain of the respective hormone receptors. The final concentrations of the donor and acceptor beads were 0.08 mg/ml and 0.02 mg/ml, respectively. Hormone concentrations from approximately 500 nM to 0 nM and receptor concentrations from 500 nM to 0 nM were cross titrated to observe expected binding patterns. Alpha buffer, the purified hormone, and the acceptor beads were added initially and allowed to incubate at room temperature for 1 hour. The donor beads were then added and the reaction was allowed to equilibrate for 1 hour at room temperature before reading the results on a Tecan Infinite M1000 Pro using the AlphaLISA filter with an excitation time of 100 ms, an integration time of 300 ms, and a settle time of 20 ms after 10 min of incubation inside the instrument. Prism 9 (GraphPad) was used to plot the data.

### Biolayer Interferometry (BLI)

Hormone binding to the extracellular domain of its receptor was tested by biolayer interferometry (BLI). Ni-NTA sensor tips were incubated in water for a least 10 min and immobilization buffer (TBS-T pH 7.6 with 1 mg/ml BSA, filtered with a 0.2 μm syringe filter (CELLTREAT 229747)) for at least 10 min before use. On an Octet K2 Biolayer Interferometer, his-tagged extracellular domain of the growth hormone receptor (GHR, BioVision 7477-10) was immobilized on the Ni-NTA sensor tip by dipping the tips in a 2.5 μg/ml solution of GHR for 400 s. Binding was assessed with an estimated growth hormone (GH) concentration ranging from 84 nM to 11.5 nM using a 40 s equilibration step (tips in buffer), a 20 s baseline step, 120 s of association (tips in the GH solution), and 900 s of dissociation. No dissociation was observed, so kinetic model curves could not be fit to the data. A commercially produced GH positive control (Tonbo Biosciences 21-7147-U010) ranging from 194 nM to 7.88 nM was also observed to have no dissociation. On a BLITZ Bio-layer interferometer, his-tagged 25 μg/ml GLP-1 receptor (GLP-1 R MyBioSource.com MBS949592) and 50 μg/ml parathyroid hormone receptor (PTH, R&D systems 5709-PR-50) were separately immobilized on Ni-NTA tips for at least 5 min. GLP-1 binding was assessed by equilibrating the tips in buffer for 30 s, measuring association by dipping tips in undiluted GLP-1 purified from CFPS reactions as described above and 100 μg/ml of positive control commercially produced GLP-1 (PeproTech 130-08) for 120 s and measuring dissociation for 120 s. PTH binding was assessed by equilibrating tips in buffer for 30 s, measuring association by dipping tips in undiluted PTH produced in CFPS and purified as previously described or in 67 μg/ml of a commercially produced positive control PTH (OriGene SA6052) for 180 s, and measuring dissociation for 120 s.

### Fluorotect SDS-PAGE Gels

Incorporation of a fluorescent lysine residue during *in vitro* translation reactions was used to test expression of SUMO-hormone constructs. CFPS reactions were run as described above with the addition of 1 μl of FluoroTect™ reagent (Promega L5001) per 50 μl CFPS reaction. After CFPS reaction completion, 3 μl of a 1:50 dilution of RNase A is added to each 50 μl reaction (Omega AC118) and incubated at 37 °C for 10 min. Samples are prepared for SDS-PAGE by adding 5 μl of sample to 4 μl of tricine sample buffer (Bio-Rad 1610739) and 1 μl of 1 M DTT and heat denaturing at 70 °C for 3 min. The samples are run on an 16.5% MINIPROTEAN^®^ Tris-Tricine gel (Bio-Rad 4563066) for 100 V for 90 min at constant voltage. The gels were imaged on a Li-COR Odyssey Fc imager on the 700 nm channel with 30 s of exposure.

### Growth Inhibition Assay

Antimicrobial peptide activity was tested with a plate-based growth inhibition study. A 5 ml culture of *Escherichia coli* MG1655 was grown overnight on Luria Broth (LB) from a glycerol stock. On the day of the assay, the overnight culture was diluted to an OD of 0.08 in LB and then grown to mid log phase (OD ~0.5). The mid log phase cells were diluted to 10^4^ cells/ml (assuming 1 OD = 8×10^8^ cells/ml) into LB with 0.005% Antifoam 204 (Sigma A6426) and 1 mg/ml BSA (Sigma-Aldrich A2153-100G). 5 μl of these cells were added to 10 μl of purified antimicrobial peptide in a UV-sterilized 384 well, clear bottom plate (Greiner Bio-One 781096). Negative controls included 10 μl of the wash buffer the hormones were dissolved in and 10 μl of a no DNA control where no DNA template was added to the CFPS reactions, but the sample was treated exactly like the other antimicrobial peptide samples. Positive controls included a final kanamycin concentration of 50 μg/ml and adding 10 μl of the LB with 0.005% Antifoam 204 and 1 mg/ml BSA without *E. coli* cells. The plate was sealed with a microseal B adhesive (Bio-Rad MSB1001) and incubated in a plate reader with continuous shaking for 18 hr at 37 °C and absorbance measurements at 600 nm every 15 min.

## Acknowledgements

This work was supported by the Defense Threat Reduction Agency (HDTRA1-21-1-0038) and the National Science Foundation (MCB 1936789). This work made use of the IMSERC MS facility at Northwestern University, which has received support from the Soft and Hybrid Nanotechnology Experimental (SHyNE) Resource (NSF ECCS-2025633), and Northwestern University. This work also used the Keck Biophysics Facility, a shared resource of the Robert H. Lurie Comprehensive Cancer Center of Northwestern University supported in part by the NCI Cancer Center Support Grant #P30 CA060553. We would like to thank Dr. Benjamin Owen and Dr. Fernando Tobias for training and assistance with mass spectrometry and Dr. Arabela Grigorescu for training and assistance with biolayer interferometry.

## Author Contributions

M.A.D., A.H.T., W.K., and M.C.J. conceived and designed experiments, M.A.D., A.H.T., and L.G. performed experiments and analyzed data, M.A.D. wrote the manuscript draft, M.A.D., A.H.T., L. G., W.K., A.S.K., and M.C.J. reviewed and edited the manuscript, and M.C.J. supervised and provided funding. All authors have read and agreed to the final manuscript.

## Conflict of Interest

M.C.J. has a financial interest in SwiftScale Biologics, Gauntlet Bio, Pearl Bio, Inc., Design Pharmaceutics, and Stemloop Inc. M.C.J.’s interests are reviewed and managed by Northwestern University in accordance with their competing interest policies. M.A.D., A.H.T., W.K., and M.C.J., have filed a provisional patent U.S. Patent Application Serial No. 63/261,045 based on the innovations described. All other authors declare no competing interests.

## Notes

### Summary of Updates

Wording has been clarified throughout, author affiliations updated, author contributions added

