## Supplementary Information for "Point-of-care peptide hormone production enabled by cell-free protein synthesis"

Supplementary Tables 1-2

Supplementary Figures 1-6

Supplementary References 1-22

### Supplementary Tables

**Table 1: A summary of the peptide hormones tested in this work.**

| Peptide Hormone | Abbreviation | Physiological Purpose |
|---|---|---|
| **Insulin Aspart** | Ins Asp | Short-acting insulin to lower blood glucose*(1)* |
| **Insulin Lispro** | Ins Lis | Short-acting insulin*(2)* |
| **Insulin Glargine** | Ins Glarg | Long-acting insulin*(3)* |
| **Regular Insulin** | Reg Ins | Short-acting insulin*(4)* |
| **Insulin A Chain** | InsA | A chain of insulin*(4)* |
| **Insulin B Chain** | InsB | B chain of insulin*(4)* |
| **Insulin A Chain Heterodimer** | Insulin A H | A chain of insulin with a leucine zipper heterodimer*(4,5)* |
| **Insulin B Chain Heterodimer** | Insulin B H | B chain of insulin with a leucine zipper heterodimer*(4,5)* |
| **Oxytocin** | Oxy | Stimulates uterine contractions*(6)* |
| **Glucagon** | Glcg | Increases blood glucose*(7)* |
| **Glucagon like peptide-1 mutant** | GLP-1 mut | A mutant of GLP-1 meant to mimic liraglutide*(8)* for diabetes and obesity treatment*(9-11)* |
| **Glucagon like peptide-1** | GLP-1 | Stimulates insulin secretion, decreases glucagon secretion, slows gastric emptying*(8,11,12)* |
| **Insulin like growth factor-1** | IGF-1 | Promotes growth of bones and tissues*(13)* |
| **Growth hormone** | GH | Promotes growth of bones and tissues*(14)* |
| **Leptin** | Lept | Regulates hunger to maintain energy homeostasis*(15,16)* |
| **Vasopressin** | Vspn | Increases blood pressure, stimulates water retention in kidneys*(17)* |
| **Angiotensin II** | Ang II | Vasoconstriction to increase blood pressure*(18)* |
| **Parathyroid hormone** | PTH | Increases serum calcium, stimulates bone formation*(19)* |
| **Somatostatin** | SST | Inhibits insulin, glucagon, GH, and thyroid stimulating hormone secretion*(20)* |
| **Leuprolide** | Leu | Gonadotropin releasing hormone that can suppress sex steroid hormone production with extended administration*(21)* |

**Table 2: Amino acid sequences of peptide hormones.**

| Name | Amino Acid Sequence |
|---|---|
| **SUMO** | MASGSDSEVNQEAKPEVKPEVKPETHINLKVSDGSSEIFFKIKKTTPLR RLMEAFAKRQGKEMDSLRFLYDGIRIQADQTPEDLDMEDNDIIEAHRE QIGG |
| **Insulin Aspartate** | FVNQHLCGSHLVEALYLVCGERGFFYTDKTRREAEDLQVGQVELGGGP GAGSLQPLALEGSLQKRGIVEQCCTSICSLYQLENYCN |
| **Insulin Lispro** | FVNQHLCGSHLVEALYLVCGERGFFYTKPTRREAEDLQVGQVELGGGP GAGSLQPLALEGSLQKRGIVEQCCTSICSLYQLENYCN |
| **Insulin Glargine** | FVNQHLCGSHLVEALYLVCGERGFFYTPKTRREAEDLQVGQVELGGGP GAGSLQPLALEGSLQKRGIVEQCCTSICSLYQLENYCG |
| **Regular Insulin** | FVNQHLCGSHLVEALYLVCGERGFFYTPKTRREAEDLQVGQVELGGGP GAGSLQPLALEGSLQKRGIVEQCCTSICSLYQLENYCN |
| **Insulin A Chain** | GIVEQCCTSICSLYQLENYCN |
| **Insulin B Chain** | FVNQHLCGSHLVEALYLVCGERGFFYTPKT |
| **A chain heterodimer** | GTKEDILERQRKIIERAQEIHRRQQEILEELERIIRKPGSSEEAMKRMLKLL EESLRLLKELLELSEESAQLLYEQR |
| **B chain heterodimer** | GTEKRLLEEAERAHREQKEIIKKAQELHRRLEEIVRQSGSSEEAKKEAKK ILEEIRELSKRSLELLREILYLSQEQKGSLVPR |
| **Oxytocin** | CYIQNCPLG |
| **Glucagon** | HSQGTFTSDYSKYLDSRRAQDFVQWLMNT |
| **GLP-1 mut** | HAEGTFTSDVSSYLEGQAAKEEFIAWLVRGRG |
| **GLP-1** | HAEGTFTSDVSSYLEGQAAKEFIAWLVKGRG |
| **IGF-1** | GPETLCGAELVDALQFVCGDRGFYFNKPTGYGSSSRRAPQTGIVDEC CFRSCDLRRLEMYCAPLKPAKSA |
| **Growth Hormone** | FPTIPLSRLFDNAMLRAHRLHQLAFDTYQEFEEAYIPKEQKYSFLQNPQ TSLCFSESIPTPSNREETQQKSNLELLRISLLLIQSWLEPVQFLRSVFAN SLVYGASDSNVYDLLKDLEEGIQTLMGRLEDGSPRTGQIFKQTYSKFD TNSHNDDALLKNYGLLYCFRKDMDKVETFLRIVQCRSVEGSCGF |
| **Leptin** | VPIQKVQDDTKTLIKTIVTRINDISHTQSVSSKQKVTGLDFIPGLHPILTLS KMDQTLAVYQQILTSMPSRNVIQISNDLENLRDLLHVLAFSKSCHLPWA SGLETLDSLGGVLEASGYSTEVVALSRLQGSLQDMLWQLDLSPGC |
| **Vasopressin** | CYFQNCPRG |
| **Angiotensin II** | DRVYIHPF |
| **Parathyroid Hormone** | SVSEIQLMHNLGKHLNSMERVEWLRKKLQDVHNFVALGAPLAPRDA GSQRPRKKEDNVLVESHEKSLGEADKADVNVLTKAKSQ |
| **Somatostatin** | SANSNPAMAPRERKAGCKNFFWKTFTSC |
| **Leuprolide** | QHWSYGLRPG |
| **Casocidin-II** | TKLTEEEKNRLNFLKKISQRYQKFALPQYLK |
| **Cecropin B** | KWKIFKKIEKVGRNIRNGIIKAGPAVAVLGEAKAL |
| **Melittin** | GIGAVLKVLTTGLPALISWIKRKRQQ |
| **Bactericidin B-2** | WNPFKELERAGQRVRDAVISAAPAVATVGQAAAIARG |
| **Catestatin** | SSMKLSFRARAYGFRGPGPQL |
| **Thrombocidin-1 D42K** | AELRCMCIKTTSGIHPKNIQSLEVIGKGTHCNQVEVIATLKKGRKIC LDPDAPRIKKIVQKKLAGDES |
| **Thrombocidin-1** | AELRCMCIKTTSGIHPKNIQSLEVIGKGTHCNQVEVIATLKDGRKICLDP DAPRIKKIVQKKLAGDES |
| **CAT-enhancer** | MEKKI *(22)* |

#### Supplementary Figures

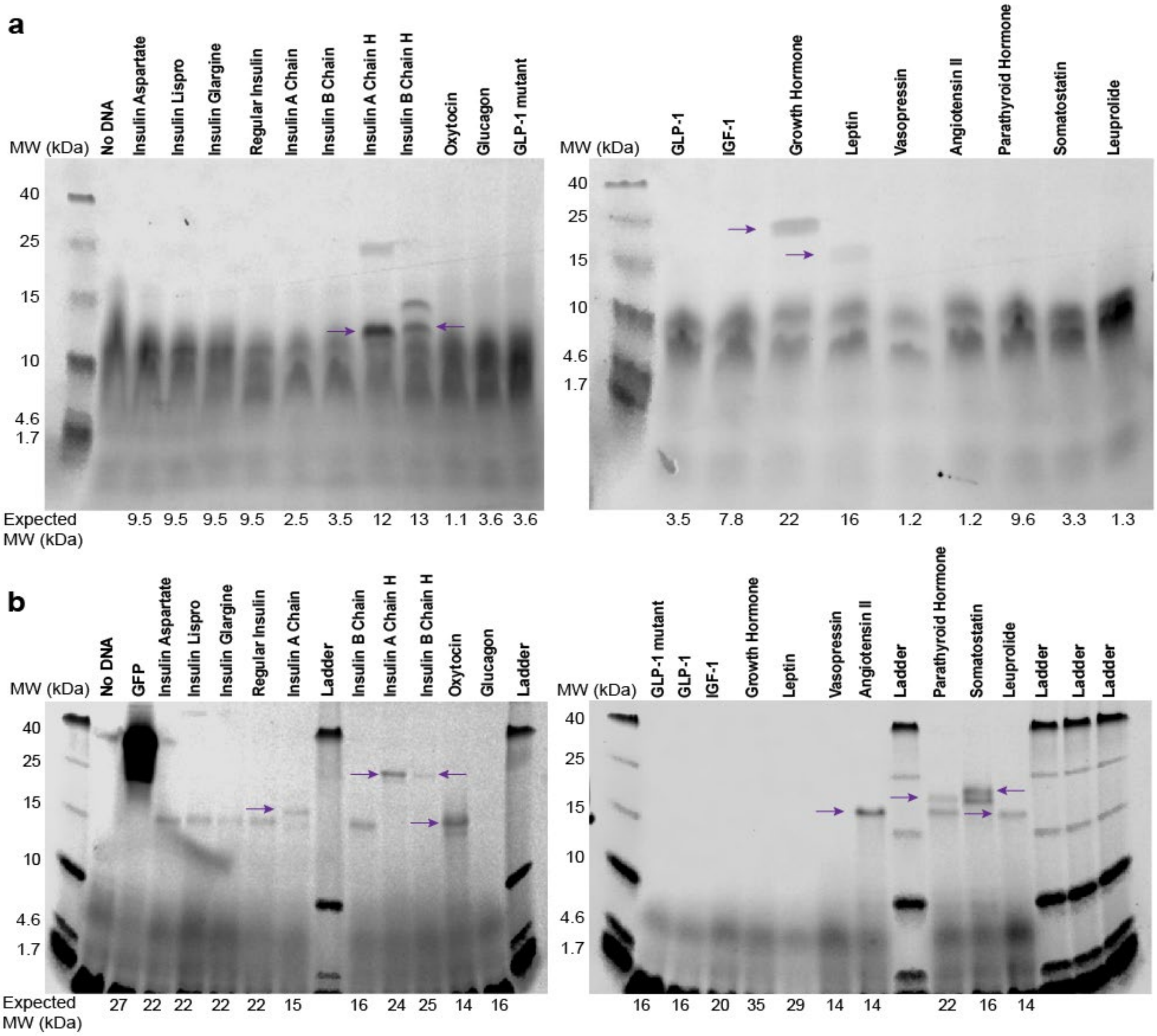


**Figure S1: SUMO fusions improve expression over untagged hormones.** (a) A panel of tagless peptide hormones (purple arrows) was screened for expression in crude BL21 extract. Incorporation of Fluorotect, a reagent with fluorescently labeled lysine residues, into the cell-free produced hormones was detected by fluorescent imaging of an SDS-PAGE gel. (b) Fluorotect incorporation was similarly used to screen expression of SUMO-hormone fusions (purple arrows) in crude BL21 extract. More full-length hormones were detected when fused with SUMO.

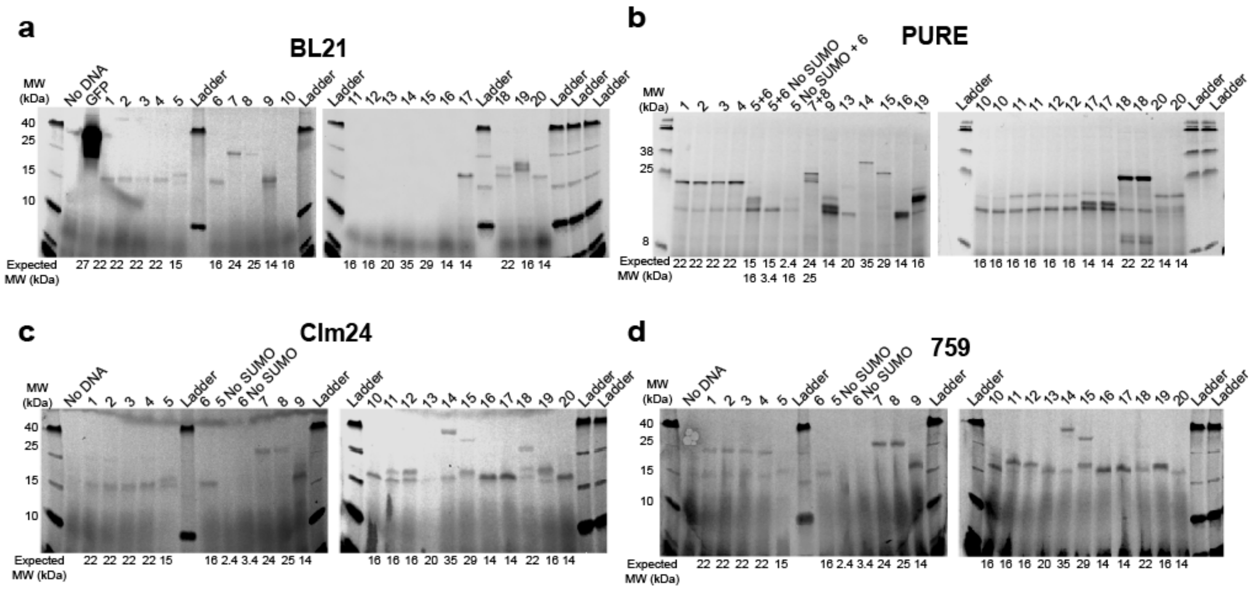


**e**

| | 1 | 2 | 3 | 4 | 5 | 6 | 7 | 8 | 9 | 10 |
|---|---|---|---|---|---|---|---|---|---|---|
| | Insulin Aspartate | Insulin Lispro | Insulin Glargine | Regular Insulin | Insulin A Chain | Insulin B Chain | Insulin A Chain Heterodimer | Insulin B Chain Heterodimer | Oxytocin | Glucagon |
| PURE | ✓ | ✓ | ✓ | ✓ | ✓ | ✓ | ✓ | ✓ | ✓ | ✓ |
| Clm24 | ✗ | ✗ | ✗ | ✗ | ✓ | ✗ | ✓ | ✓ | ✓ | ✓ |
| BL21 | ✗ | ✗ | ✗ | ✗ | ✓ | ✗ | ✓ | ✓ | ✓ | ✗ |
| 759 | ✓ | ✓ | ✓ | ✓ | ✓ | ✓ | ✓ | ✓ | ✓ | ✗ |
| | 11 | 12 | 13 | 14 | 15 | 16 | 17 | 18 | 19 | 20 |
| | Glucagon Like Peptide 1 mutant (GLP-1 mut) | Glucagon Like Peptide 1 (GLP-1) | Insulin Like Growth Factor | Growth Hormone (GH) | Leptin | Vaso-pressin | Angiotensin II | Parathyroid Hormone (PTH) | Somato-statin | Leuprolide |
| PURE | ✓ | ✓ | ✓ | ✓ | ✓ | ✓ | ✓ | ✓ | ✓ | ✓ |
| Clm24 | ✓ | ✓ | ✗ | ✓ | ✓ | ✓ | ✓ | ✓ | ✓ | ✓ |
| BL21 | ✗ | ✗ | ✗ | ✗ | ✗ | ✗ | ✓ | ✓ | ✓ | ✓ |
| 759 | ✓ | ✓ | ✗ | ✓ | ✓ | ✓ | ✓ | ✗ | ✓ | ✓ |

**Figure S2: Screen of a panel of peptide hormones in lysates from different *E. coli* strains for improved expression.** (a) A panel of peptide hormones was screened for expression in BL21. The same figure from Fig S1b is repeated here for comparison. Incorporation of Fluorotect, a reagent with fluorescently labeled lysine residues, into the cell-free produced hormones was detected by fluorescent imaging of an SDS-PAGE gel. Lanes are labeled with sample numbers given in panel (e). Fluorotect incorporation was similarly used to screen expression in PUREfrex 2.1 (b), Clm24 (c), and 759 (d) extracts. (e) A summary of expression systems which produced full length peptide hormones is shown. Green checks indicate full length expression and red crosses indicate that full length expression was not observed.

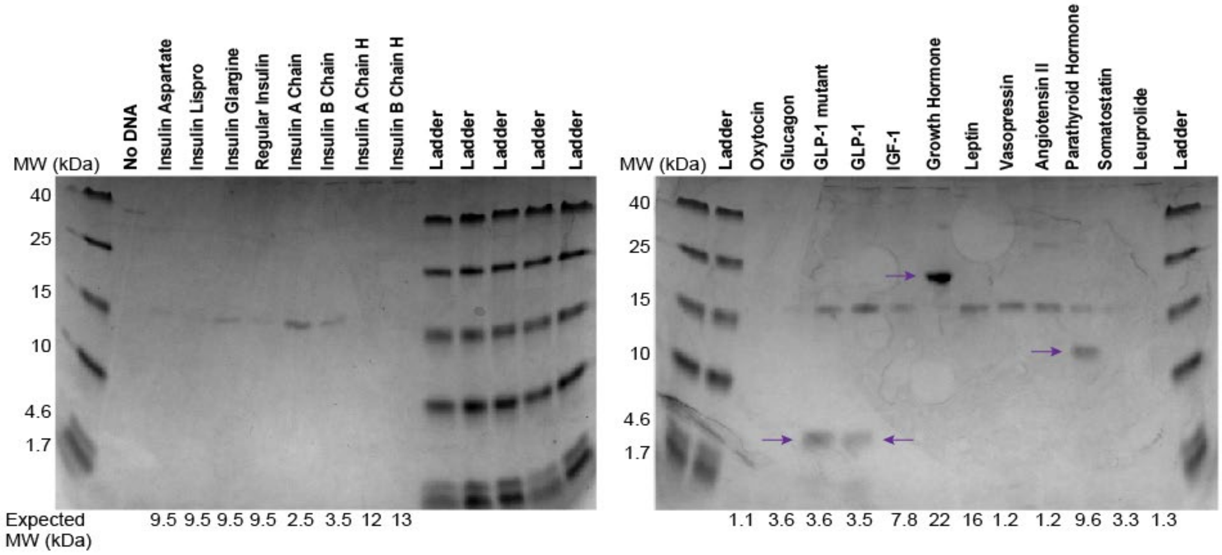


**Figure S3: Screen of the purification procedure on the full peptide hormone panel.** After expression in 759 extract and undergoing the 2-in-1 purification and cleavage procedure, an initial screen showed recovery of untagged GLP-1 mut, GLP-1, GH, and PTH (purple arrows) on a Coomassie stained SDS-PAGE gel.

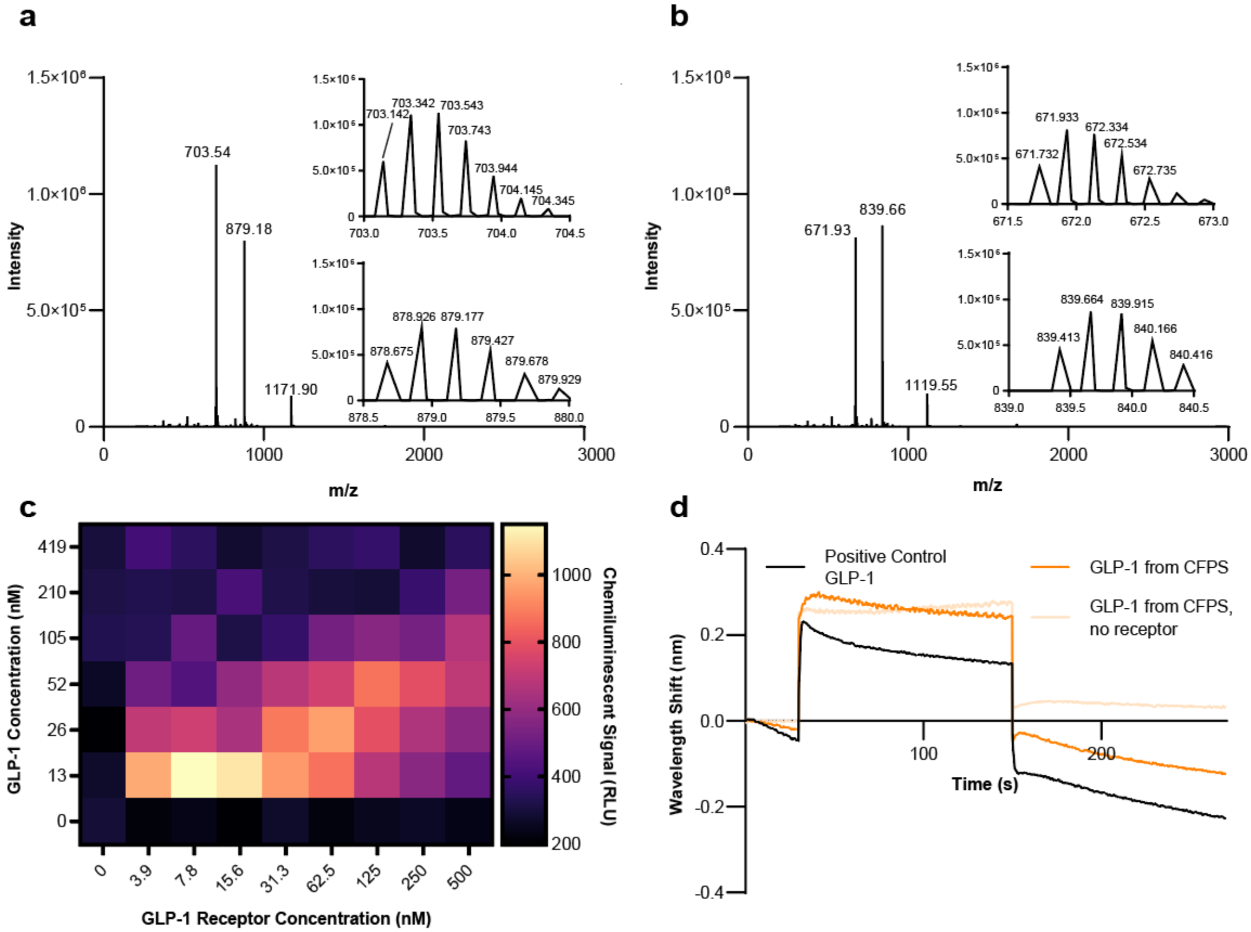


**Figure S4: Characterization of GLP-1 and GLP-1 mut by LC-MS, AlphaLISA, and BLI indicates cell-free produced hormone is intact and may be binding to its receptor.** (a) Peptide liquid chromatography-mass spectrometry (LC-MS) data shows intact GLP-1 mut is recovered after purification. The full scan is shown with insets displaying detailed isotopic distributions of the +5 charge state ions at average m/z of 703.54 m/z and the +4 charge state ions at average m/z of 879.18. The monoisotopic masses observed (3510.67 Da) match the theoretical monoisotopic mass of GLP-1 mut (3510.72 Da) with an error of 15 ppm. (b) Peptide LC-MS data shows intact glucagon-like peptide (GLP-1) is recovered after purification. The full scan is shown with insets displaying detailed isotopic distributions of the +5 charge state ions at average m/z of 671.96 m/z and the +4 charge state ions at average m/z of 839.66. The monoisotopic masses observed (3353.62 Da) match the theoretical monoisotopic mass of GLP-1 (3353.67 Da) with an error of 15 ppm. (c) GLP-1 and GLP-1 receptor concentrations were cross titrated, and binding was assessed with AlphaLISA. (d) GLP-1 binding to the extracellular domain of its receptor was assessed by BLI. No binding was observed with GLP-1 produced in CFPS or the commercially available GLP-1, likely because the hormone is too small to create a difference in optical thickness that can be detected by the instrument.

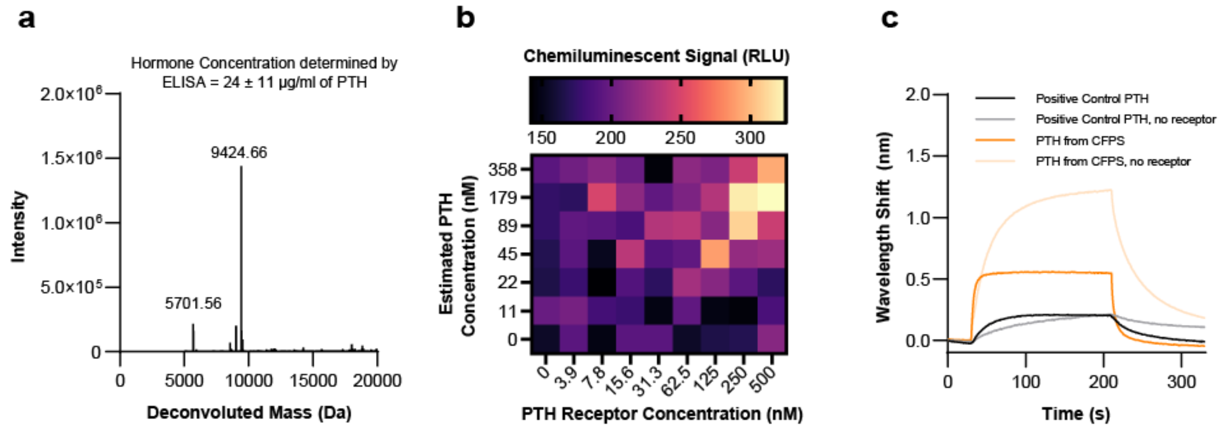


**Figure S5: Characterization of PTH by ELISA, LC-MS, AlphaLISA, and BLI, indicates hormone is intact and shows some binding with its receptor.** (a) PTH identification and quantification was determined by sandwich ELISA with 4 dilutions and intact protein LC-MS. The observed mass (9424.66 Da) on a deconvoluted spectrum matches the expected average mass of PTH (theoretical: 9424.73 Da) with an error of 7.4 ppm. (b) PTH and PTH receptor concentrations were cross titrated, and binding was assessed with AlphaLISA. (d) PTH binding to the extracellular domain of its receptor was assessed by BLI. A large background signal for both commercially produced PTH positive control and PTH produced in CFPS prevented accurate detection of binding. PTH is slightly smaller than the molecular weight cutoff for this BLI instrument.

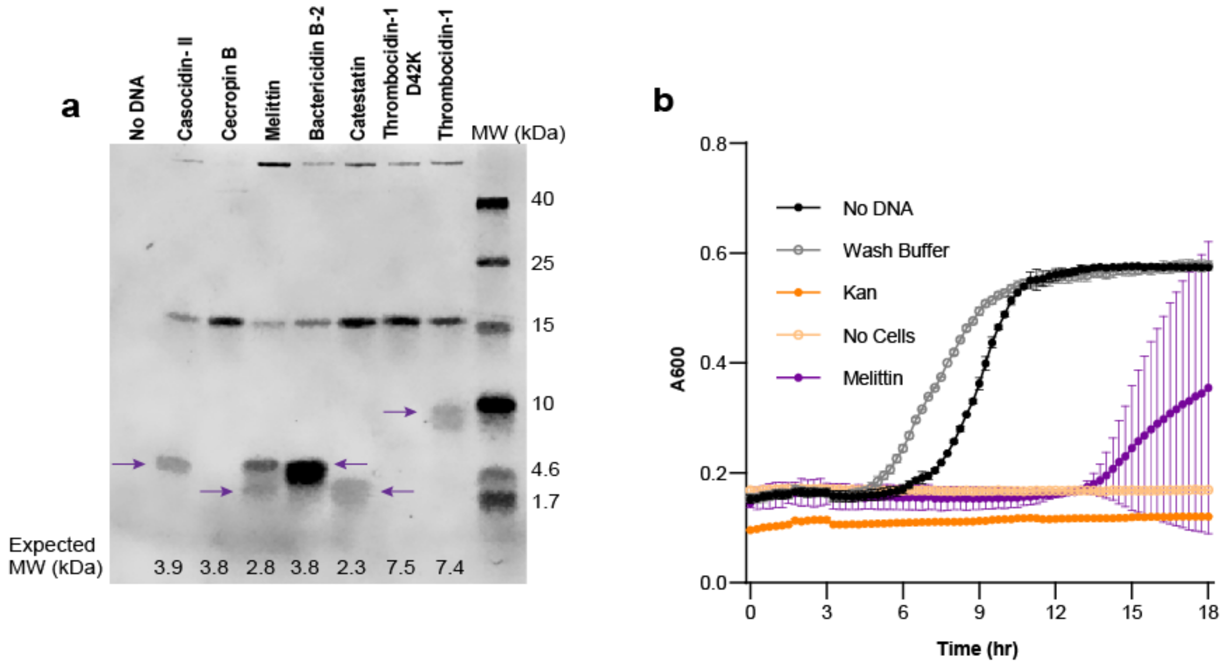


**Figure S6: Anti-microbial peptides can also be purified using the described method and are active in the prevention of growth of *E. coli* MG1655.** (a) A Coomassie SDS-PAGE gel shows recovery of anti-microbial peptides (purple arrows) after the Strep-SUMO purification procedure described in Figure 3a. (b) Melittin inhibits growth of *E. coli* MG1655 during overnight incubation in a 384 well plate. Error bars represent 1 s.d. of (n=2) replicates. Error bars not shown are smaller than the symbol.
